# Heat and humidity sensors that alert mosquitoes to nearby human hosts

**DOI:** 10.1101/2022.04.05.487046

**Authors:** Willem J. Laursen, Gonzalo Budelli, Elaine C. Chang, Rachel Gerber, Ruocong Tang, Chloe Greppi, Rebecca Albuquerque, Paul A. Garrity

## Abstract

Mosquito-borne diseases sicken >500,000,000 people annually, killing >500,000^1^. Mosquito host-seeking is guided by multiple host-associated cues, which combine to drive blood feeding in a manner that remains poorly understood^2,3^. While heat is a powerful mosquito attractant, recent studies indicate that disruption of heat seeking has little impact on host detection by the malaria vector *Anopheles gambiae*^4^, suggesting other cues act alongside heat in the complex sensory environment of a human host. Here we show mosquitoes require *Ir93a* (an Ionotropic Receptor^5^) to maintain attraction to a human host and feed on warmed blood. Using *Ir93a*, we uncover the previously uncharacterized mosquito hygrosensory system, and show *Ir93a* is required for humidity detection by humidity sensors (hygrosensors) as well as temperature detection by thermosensors, and for attraction to each stimulus. These findings indicate that hygrosensation and thermosensation function in parallel, driving host proximity detection in response to the overlapping heat and humidity gradients humans produce^6,7^. These host cue sensors appear to have arisen by co-opting existing sensors of physical cues rather than *de novo*, as *Ir93a*-dependent thermo- and hygro-sensors support physiological homeostasis in non-blood-feeding insects^8–11^. While *Ir93a* is conserved among arthropods, reliance on heat and humidity evolved independently in multiple blood-feeding lineages, suggesting multiple, vector-specific implementations of this common host-seeking strategy.

## Main Text

Blood feeding by vector mosquitoes is critical for spreading viruses and parasites responsible for devastating diseases like dengue and malaria^1^. Vector mosquitoes acquire and transmit pathogens during blood feeding, a behavior through which they also obtain nutrients needed to reproduce. These multiple roles allow modest reductions in blood feeding to significantly reduce disease burden, with models suggesting a two-fold decrease in blood feeding rate can elicit an ~8-fold drop in the reproductive ratio (R value) of a vector-borne disease^12^. This behavior is driven by multiple host-associated cues: visual, chemical, and physical^2,3,13^. Visual and chemical cues can impact mosquito behavior from meters away, but physical cues like temperature are encountered near the host and promote host-proximal behaviors like landing and probing^2,3^. In the malaria vector *Anopheles gambiae,* the Ionotropic Receptor (IR) *Ir21a* was recently shown to mediate detection of host-associated temperatures^4^. But while loss of *Ir21a* impaired heat seeking, the more ethologically relevant behavior, attraction to a human hand, was largely unaffected^4^. We hypothesized this might reflect the influence of a parallel cue emanating from the host.

In addition to short-range thermal gradients, humans also generate short-range humidity gradients. The two gradients are largely coextensive and form a “boundary layer” of heated, moistened air that surrounds the human body^6,7^, making humidity a candidate for this parallel cue (Fig. 1a, b). While humidity sensation (hygrosensation) promotes host seeking^14^, in contrast to other sensory modalities implicated in host seeking, such as vision, olfaction, taste and thermosensing, the neurons involved in mosquito hygrosensing have not been established^15^, and no molecular receptors described. In *Drosophila*, hygrosensation depends on multiple receptors that, like *Ir21a*, belong to the IR family: hygrosensors activated by moist air (Moist Cells) require *Ir68a* and dry air (Dry Cells) require *Ir40a*^9–11,16^. Both hygrosensors also require *Ir93a*, an IR co-receptor that acts alongside stimulus-specific IRs like *Ir68a* and *Ir40a*^9–11^. Importantly, *Drosophila Ir93a* also acts with the fly ortholog of *Ir21a* to mediate thermosensing^8^. While *Drosophila* do not blood feed and diverged from mosquitoes ~250 million years ago^17^, this raised the possibility that the mosquito ortholog of *Ir93a* might provide a single gene target whose inactivation could cripple multiple systems involved in sensing host proximity, potentially overcoming functional redundancies to interfere with host proximity detection.

**Fig. 1.**
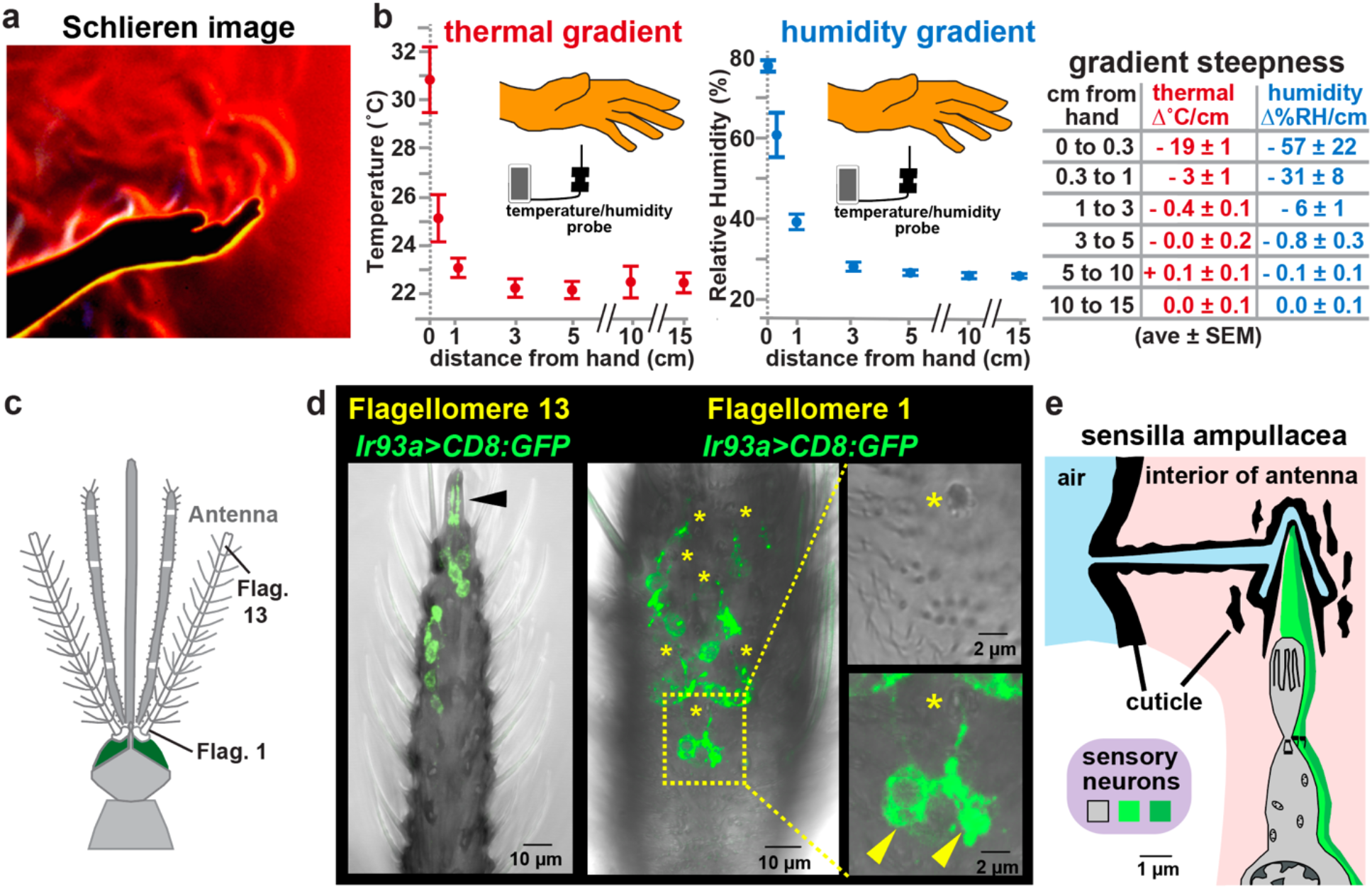
Humans generate co-extensive thermal and humidity gradients, and mosquitoes express *Ir93a>CD8:GFP* in potential thermo- and hygro-sensors. **(a)** Refractive index variations near hand, reflecting temperature and humidity. Yellow/white, inner boundary layer. Image, G. Settles; for details^7^. (**b)** Gradients beneath hand. n=6. **(c)** Mosquito anterior, after^30^. **(d)** *Ir93a^pore^>CD8:GFP*. Insets (right) indicate sensilla ampullacea (top), with CD8:GFP overlay (bottom). Black arrowhead, coeloconic sensilla. Asterisks, sensilla ampullacea. Yellow arrowheads, cell bodies. **(e)** Sensilla ampullacea, after^15^.

*Anopheles gambiae Ir93a* was disrupted by CRISPR/Cas9-aided integration of a knock-in cassette in exonic locations essential for IR activity: the putative ion pore (creating *Ir93a^pore^*) and the second transmembrane domain (creating *Ir93a^TM2^*) (Extended Data Fig. 1). Both *Ir93a^pore^* and *Ir93a^TM2^* were homozygous viable and express the QF2 transcriptional activator under endogenous *Ir93a* regulatory element control. Expression was characterized in *Ir93a^pore^>CD8:GFP/Ir93a^wt^* and *Ir93a^TM2^*>*CD8:GFP/Ir93a^wt^* females (collectively termed *Ir93a>CD8:GFP*), which express QF2-responsive CD8:GFP in similar patterns and contain one wild type *Ir93a* allele.

*Ir93a>CD8:GFP* was expressed in the antenna (Fig 1. c, d; Extended Data Fig. 2), but not other tissues implicated in host detection, like the maxillary palps and labellum, consistent with RNA-seq studies^18^. Expression was detected in all flagellomeres (Extended Data Fig. 2a), and included neurons innervating coeloconic sensilla in flagellomere 13, which contain thermosensory neurons (Fig. 1d), as well as ~4 neurons per flagellomere which innervate trichoid sensilla and co-express the olfactory co-receptor Orco (Extended Data Fig. 2b, d, e). Expression was also detected in pairs of neurons innervating a class of ~9 functionally uncharacterized sensilla in flagellomeres 1 and 2, the sensilla ampullacea (Fig. 1d, Extended Data Fig. 2c). These “peg-in-tube” sensilla reside inside the antenna, at the base of a tube opening at the antennal surface (Fig. 1e)^15^. Their location and lack of pores for chemical entry has led to their classification as thermosensory^15^. However, hygrosensory neurons in other insects innervate poreless sensilla^19^, suggesting they could potentially serve this role.

The humidity responsiveness of sensilla ampullacea-associated neurons was examined by trans-cuticular calcium imaging (Fig. 2a-e). In control animals (containing one wild type *Ir93a* allele), the pairs of *Ir93a>GCaMP6f*-labeled neurons showed opponent responses: one neuron, the Moist Cell, was activated by humid air, and the other, the Dry Cell, was activated by dry air (Fig. 2a-e). In *Ir93a* mutants, both responses were dramatically reduced (Fig. 2a-e), demonstrating that sensilla ampullacea harbor *Ir93a-*dependent Moist and Dry Cell hygrosensors.

**Fig. 2:**
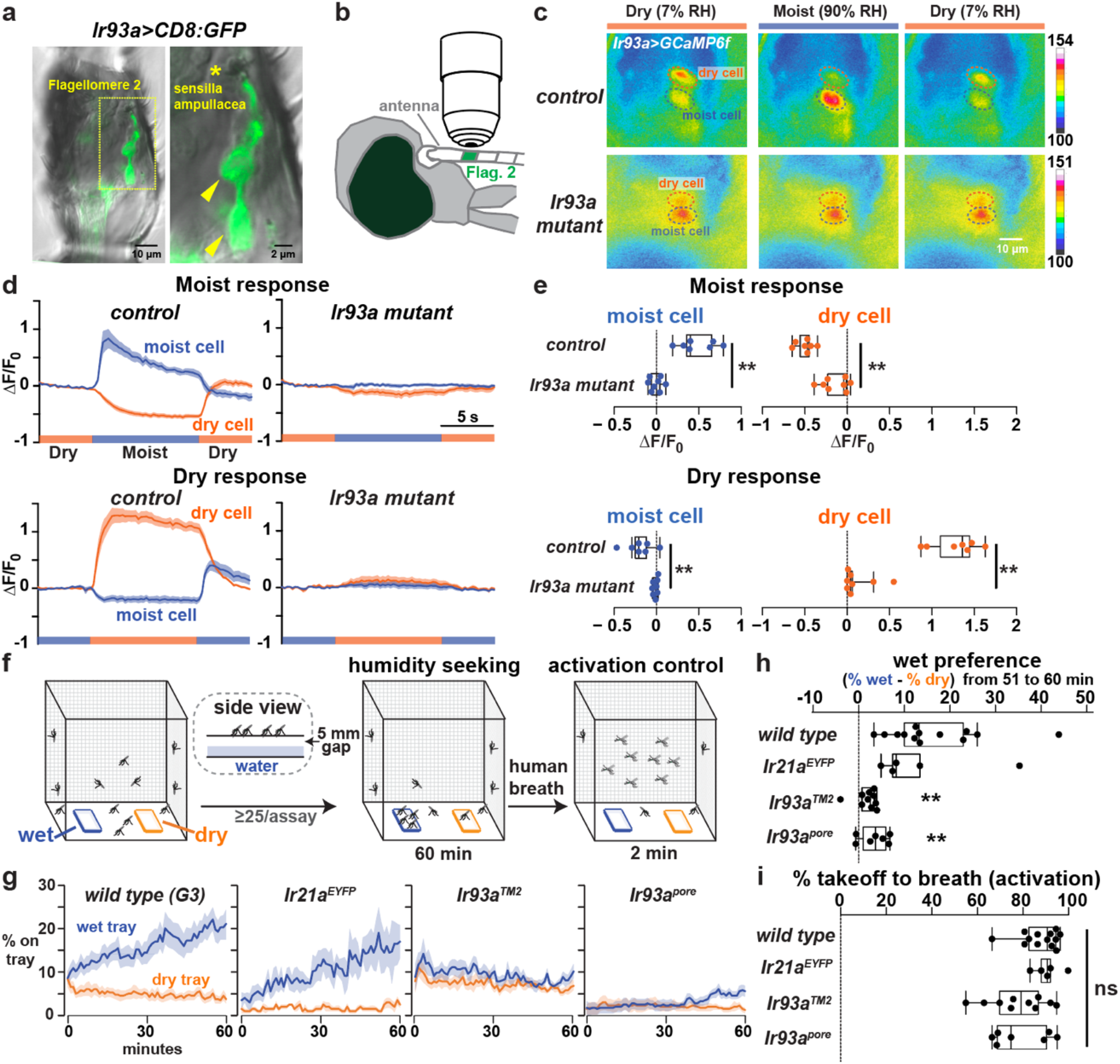
*Ir93a* mediates hygrosensation and humidity seeking. **(a)** *Ir93a^Pore^>CD8:GFP-* neurons in flagellomere 2. Asterisk, sensilla ampullacea. Yellow arrowheads, cell bodies**. (b)** Trans-cuticular imaging. **(c)** *Ir93a^pore^>GCaMP6f* fluorescence at indicated relative humidity (RH). Dashed ovals, soma. **(d)** Moist (90% RH) and dry (7% RH) responses. Mean +/-SEM. Control *(Ir93a^pore^/+*), n=7 animals. *Ir93a* mutant (*Ir93a^pore^/Ir93a^pore^*), n=8. F_0_ = average 5.5s to 1s pre-stimulus-switch. **(e)** Fluorescence change upon switching between moist and dry. ΔF = F(average 3s to 5.5s post-switch) -F_0_. ** p<0.001, t-test. **(f)** Humidity-seeking assay. **(g)** Mosquitoes landed on trays. Mean +/- SEM. 25-73 females/assay. *wt*, n=13 assays. *Ir21a^EYFP^*, n=5. *Ir93a^TM2^*, n=10. *Ir93a^pore^*, n=7. **(h)** Average (% on wet - % on dry) 51 to 60 min. ** p<0.01 versus *wt*, Steel with control. **(i)** Percent taking flight within 2 minutes after breath exposure.

To test *Ir93a’s* contribution to humidity-seeking behavior, mated, non-blood fed females were provided a choice between two trays, one dry and one water-containing; both were mesh-covered, preventing liquid contact, but permitting water vapor escape (Fig. 2f). Wild type preferentially accumulated on the water-containing tray, but this preference was strongly reduced in *Ir93a* mutants (Fig. 2g, h). In contrast, mosquitoes lacking *Ir21a*, which mediates heat seeking^4^, showed no significant decrement (Fig. 2g, h). All genotypes took flight similarly in response to human breath at assay’s end (Fig. 2I), confirming the sensory specificity of the *Ir93a* mutants’ defect.

To test whether *Ir93a* also mediates thermosensing, we examined *Ir21a*-dependent Cooling Cells at the antenna’s tip (Fig. 3a)^4^. In *Drosophila*, Cooling Cells require both *Ir21a* and *Ir93a* to sense temperature^8^, suggesting *Anopheles Ir93a* might act similarly. Cooling Cells exhibit baseline spiking at constant temperature, and their spike rates transiently rise upon cooling and transiently fall upon warming (Fig. 3b, c)^4^. Cooling Cell thermosensitivity was abolished in *Ir93a* mutants (Fig. 3b, c), confirming its thermosensory function.

**Fig. 3:**
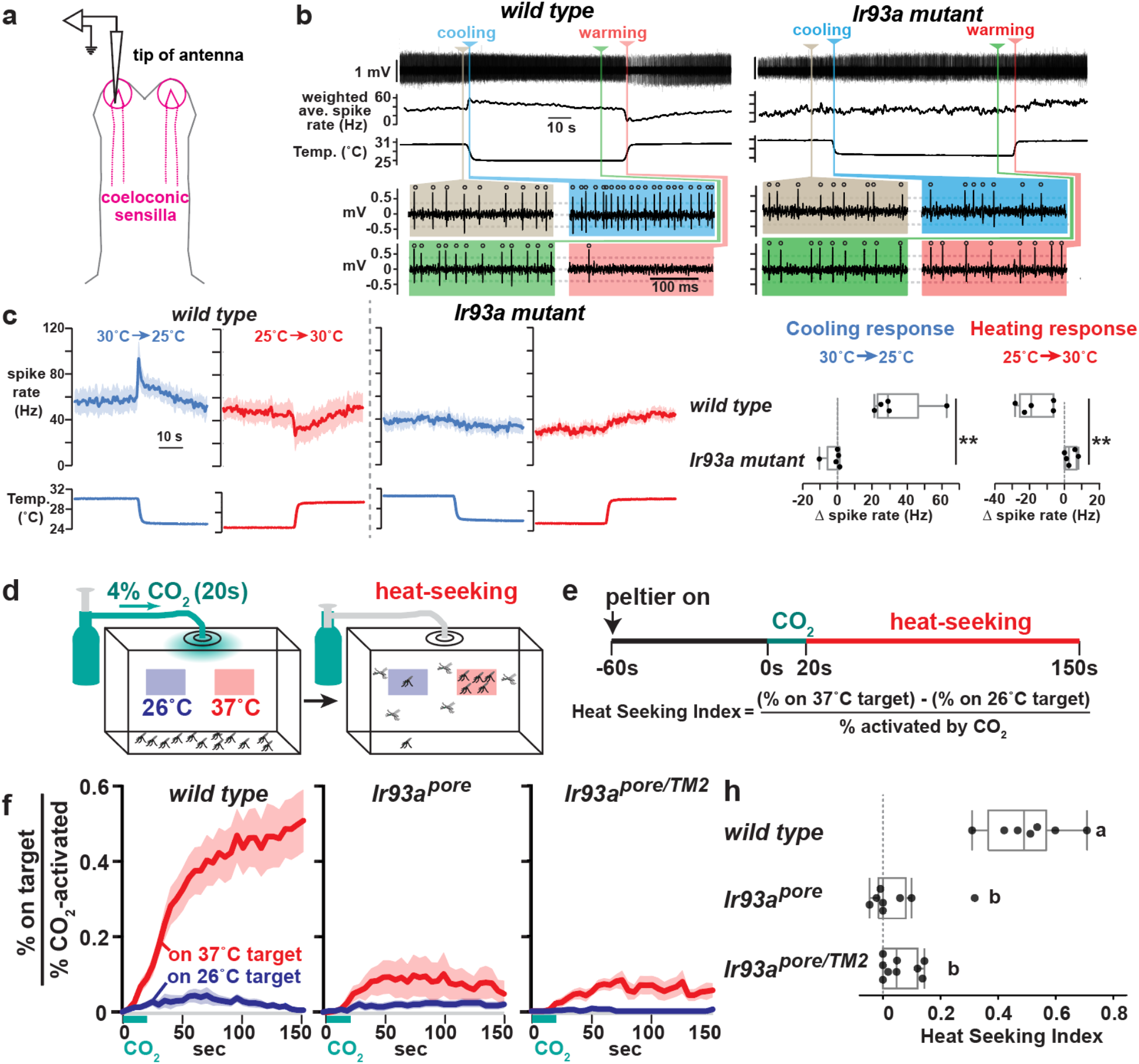
*Ir93a* is required for thermosensation and heat seeking. **(a)** Coeloconic sensilla recording. **(b)** *wild type (G3)* and *Ir93a* mutant (*Ir93a^TM2^*) recordings. Weighted average spike rates (1s triangular window). Colored panels highlight activity at indicated times. Open circles, spikes. Dashed lines, thresholds. **(c)** Peri-stimulus time histograms (average +/-SEM). Cooling, blue. Warming, red. *wild type*, n=5 animals. *Ir93a^TM2^*, n=5. Cooling response = (average Hz 0.2s- 0.7s after cooling onset) – (average Hz 5s-10s pre-cooling). ** p=0.009, Wilcoxon. Heating response = (average Hz 0.5s-1.5s after heating onset) – (average Hz 5s-10s pre-heating). ** p= 0.0026, t-test. **(d)** Heat seeking assay. **(e)** Assay time-course and index calculation. **(f)** Fraction of CO_2_-activated mosquitoes landed on targets. Average +/- SEM. Wild type, n=7 assays. *Ir93a^pore^/Ir93a^pore^*, n=7. *Ir93a^pore^/Ir93a^TM2^*, n=9. 46-50 females/assay. **(g)** Heat seeking index, average 130s-150s. Letters denote distinct groups, p<0.01, Steel-Dwass.

*Ir93a”s* contribution to heat seeking behavior was also tested. Exposure to 4% CO_2_ for 20 sec (simulating breath) prompted mosquitoes to take flight (Extended Data Fig. 3), and, in wild type, ~50% of these “activated” animals landed on a 37°C target, largely ignoring an adjacent 26°C target (Fig. 3d-g). *Ir93a* was important for heat seeking, as only ~10% of activated *Ir93a* mutants landed on the 37°C target (Fig. 3d-g), a defect resembling *Ir21a* mutants^4^.

Having established *Ir93a’* s importance for detecting heat and humidity presented in isolation, *Ir93a’* s function was examined in the more complex sensory environment near a human host. Females were placed in a mesh cage, activated by five human breaths, and presented a human hand ~3 mm above the cage (Fig. 4a). This gap prevents physical contact with the host, but still provides exposure to host-generated chemical, visual, thermal and hygrosensory cues. ~70% of wild type mosquitoes approached and landed below the hand, sustaining this attraction for many minutes (Fig. 4b-e, Movie 1). When the hand was removed, they rapidly departed (Fig. 4b; Movie 1), indicating their continued presence depended on detecting cues from the hand.

**Fig. 4:**
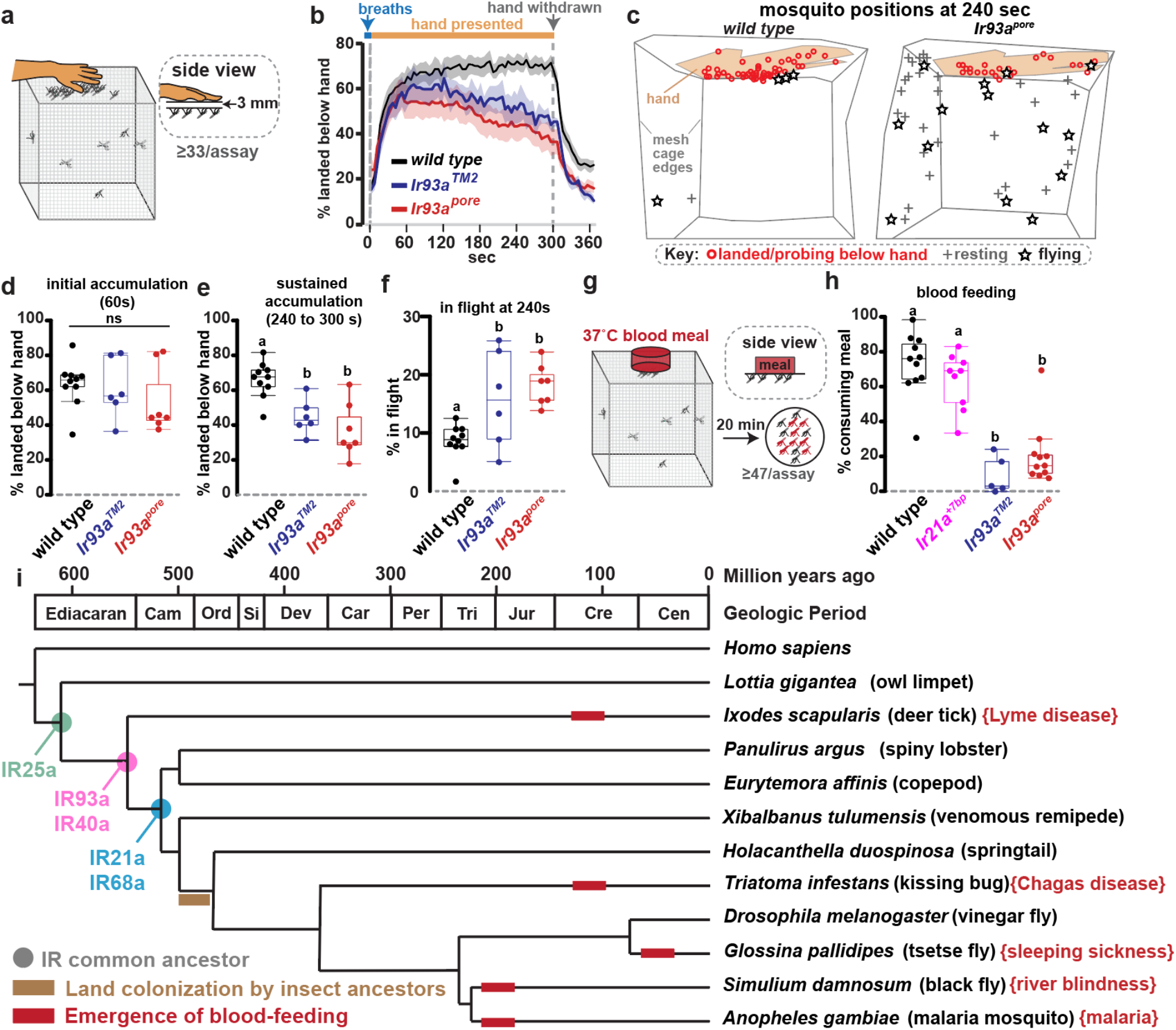
*Ir93a* is required to maintain close-range host attraction and promote blood feeding and is conserved among arthropods. **(a)** Host attraction assay. **(b)** Average +/-SEM. 41-76 females/assay. *wild type*, n=10 assays. *Ir93a^TM2^*, n=6. *Ir93a^pore^*, n=7. **(c)** Tracings of mosquito locations and behaviors from assay videos 240s post-hand presentation. (**d-e)** Mosquitoes landed under hand 60s (ANOVA, p=0.851) (**d**) and 240-300s post-hand presentation (letters denote distinct groups, Tukey HSD, alpha=0.01) (**e**). **(f)** Animals in flight 240s post-hand presentation. Tukey HSD, alpha=0.05. **(g)** Blood feeding assay, with human blood meal behind collagen membrane. **(h)** Blood consumption. 47-76 females/assay. *wild type*, n=11 assays. *Ir21a^EYFP^*, n=9. *Ir93a^TM2^*, n=5. *Ir93a^pore^*, n=11. Steel-Dwass, p<0.05. **(i)** Ionotropic Receptor distribution. Cladogram, land colonization and blood feeding emergence times based on ^17,27,31–34^. Cam, Cambrian. Ord, Ordovician. Si, Silurian. Dev, Devonian. Car, Carboniferous. Per, Permian. Tri, Triassic. Jur, Jurassic. Cre, Cretaceous. Cen, Cenozoic (era). Brackets denote a disease transmitted by each vector.

In *Ir93a* mutants, attraction to the hand resembled wild type at early time points, but steadily declined to well below wild type over the course of the assay (Fig. 4b, d, e; Movie 2). Furthermore, as illustrated in Movie 2, the decline in *Ir93a* mutant mosquitoes landed beneath the hand was frequently accompanied by a significant increase in *Ir93a* mutants engaged in sustained flights within the cage (Fig. 4c, f; Movie 2). These animals appear to remain engaged in host-seeking, despite not landing below the hand presented. This contrasts with *Ir21a* mutants, which sustain attraction to the hand normally^4^. This indicates that *Ir93a’* s role in host sensing extends beyond *Ir21a*-dependent thermosensing to the detection of additional short-range host cues. Thus, *Ir93a* is not required for initial approach in this assay, in which longer range visual and chemical cues are also present, but rather for sustaining close-range host attraction, consistent with a key role for *Ir93a* in detecting cues that signal host proximity.

As short-range interactions are vital for blood feeding, consumption of a human blood meal from a collagen membrane feeder was also examined (Fig. 4g). We previously found that when presented with two blood meals of differing temperature, wild type females strongly preferred warmer blood, and loss of *Ir21a* reduced this preference^4^. However, if presented with only a single 37°C blood meal (Fig. 4g), *Ir21a* mutant consumption did not differ significantly from wild type (Fig. 4h). This indicates that although *Ir21a-*dependent thermosensing promotes blood feeding, other pathways suffice to drive the behavior in *Ir21a’* s absence. In contrast, blood feeding was strongly reduced in *Ir93a* mutants: ~70% of wild type mosquitoes consumed the 37°C blood meal, compared to only ~10 to ~20% of *Ir93a* mutants (Fig. 4h). Thus, the loss of *Ir93a* has a more extensive impact on behavior than the loss of *Ir21a*, with *Ir93a* loss impairing both sustained host attraction and membrane blood feeding, while *Ir21a* loss impairs neither.

The behavioral deficits of the *Ir93a* mutant match *Ir93a’* s observed roles in hygrosensing and thermosensing. Human hosts as well as 37°C collagen membrane feeders provide both hygrosensory and thermosensory cues, each generating spatially overlapping temperature and humidity gradients (Fig. 1b, Extended Data Fig. 4a). Consistent with parallel contributions from both modalities to behavior, using either room temperature blood or a water vapor impermeable membrane decreased feeding from an artificial feeder, while using both abolished feeding (Extended Data Fig. 4b).

These data demonstrate that the malaria vector *Anopheles gambiae* detects heat, humidity, and nearby hosts through a battery of sensors reliant on a common receptor, *Ir93a*. In *Drosophila*, each *Ir93a* function involves an additional stimulus-specific IR: *Ir21a* in Cooling Cells^8^, *Ir40a* in Dry Cells^10,11^ and *Ir68a* in Moist Cells^9^. *A. gambiae* encodes orthologs of these IRs as well as the broadly expressed co-receptor *Ir25a*, which also contributes to these responses^8–11,20^. As *Ir93a* expression is also detected in some mosquito olfactory neurons, there could be yet more stimulus-specific IRs. In *Anopheles*, *Ir21a* mediates thermosensing^4^, while *Ir25a, Ir40a* and *Ir68a* remain to be examined. All these IRs are conserved from crustaceans to insects^20–22^, indicating they emerged ~250 million years before blood feeding (Fig. 4i). *Ir93a* and its partners would therefore have initially contributed to other functions, in particular homeostatic functions like thermoregulation and water balance, as they do in contemporary *Drosophila*^8–11,16^. This suggests that rather than the *de novo* generation of dedicated thermosensors and hygrosensors for host-seeking, the evolution of mosquito hematophagy involved co-opting existing detectors of abiotic environmental stimuli.

More broadly, *Ir93a*, as well as *Ir21a, Ir25a, Ir40a* and *Ir68a*, emerged before the ancestors of modern insects colonized land, indicating that *Anopheles* heat- and humidity-seeking involves a molecular toolkit that emerged in the Cambrian oceans (Fig. 4i). The presence of orthologs of hygrosensory IRs in aquatic creatures is striking, as hygrosensation is inherently terrestrial. However, *Ir93a-*dependent hygrosensors innervate poreless sensilla, suggesting they do not detect humidity directly, but via a humidity-dependent stimulus transmitted through the cuticle, like force^23^. Crustacean orthologs of these insect IRs might therefore detect similar sensory stimuli produced by different external cues, for example, pressure at different water depths. Finally, the absence of close mammalian relatives of IRs may enhance their attractiveness as potential targets for control agents.

*Ir93a* and its partners are conserved across blood-feeding insects (Fig. 4i). While *Ir93a’*s functions in these other animals remain unexplored, heat and humidity drive blood feeding in multiple disease vectors, including *Aedes* mosquitoes, kissing bugs, tsetse flies and stable flies^14,24–26^. This represents evolutionary convergence, as blood feeding emerged independently in mosquitoes, kissing bugs and flies^17,27^ (Fig. 4i). It will therefore be of interest to determine how broadly *Ir93a*-dependent signaling has been co-opted for host seeking in other vectors, particularly as insects possess alternative, IR-independent pathways for detecting physical cues^28^. Widespread reliance on heat and humidity as host cues may reflect their association with a wide range of potential hosts. This contrasts with host-specific cues, like volatile organic chemicals, which facilitate host discrimination^13,29^. Together, heat and humidity not only provide blood-feeding insects with spatially overlapping stimulus gradients to maximize feeding on favored hosts, their ubiquity allows them to promote feeding on non-favored hosts and help ensure this behavior, which is essential for vector mosquito reproduction, proceeds even under unfavorable conditions.

## Methods

### Measurement of humidity and temperature gradients

Temperature and humidity gradients associated with a human hand were measured under identical conditions to the host approach assay (see below). A human hand was placed on a 3mm high plastic mesh spacer situated on top of a mesh cage (Bugdorm, 17.5×17.5×17.5cm). Temperature and relative humidity levels were recorded at specific distances below the hand using a Sensirion EK-H4 sensor. Gradients were measured in triplicate from two independent volunteers in a randomized order. To calculate the steepness, gradients from each individual hand were calculated before being averaged.

### Animal care

*Anopheles gambiae* mosquitoes (G3 strain) were grown in incubators (Percival Scientific) maintained at 27°C and 70-80% relative humidity with a 12-hour light/dark cycle. Larvae were raised in trays of deionized water and fed a mixture of powdered commercial fish foods (TetraMin flakes #77101 and TetraPond sticks #16467, Tetra Co., Melle, Germany) and Koi pellets (Kaytee, #100033588). Prior to eclosion, pupae were transferred to mesh cages (Bugdorm, MegaView Science Co., Ltd., Taiwan) where they remained as adults. Except where otherwise noted, adults were provided with constant access to water and 10% w/v glucose. A collagen membrane feeding system (Hemotek, ltd., Blackburn, UK) was used to provide warmed human blood (Research Blood Components, Watertown, MA) to adult females. Feeding procedures were approved and monitored by Brandeis Institutional Biosafety Committee (protocol #16015).

### Cloning

#### AgU6-IR93a gRNA_AgVasa-Cas9 plasmids

The coding sequence of *AgIR93a* (AGAP000256) was screened for Cas9 target sites devoid of predicted off-target activity using CHOPCHOP (https://chopchop.cbu.uib.no). One site in the putative pore and one site in the second transmembrane (TM2) region were chosen to create targeted knock-ins via homology directed repair. Complimentary oligonucleotides corresponding to the pore (5’-tgctGAACTGCTTCTGGTACATCTA -3’ and 5’-aaacTAGATGTACCAGAAGCAGTTC-3’) and TM2 (5’-tgctGAATCATCATCGGTACCTGG -3’ and 5’-aaacCCAGGTACCGATGATGATTC-3’) sites were annealed and cloned into the AgVasa-Cas9-sgRNA backbone vector^4^ using Bsa1 restriction sites.

#### AgIR93a-T2A-QF2_attp_3xp3-XFP_attP donor plasmid

To generate an RFP-marked version of the previously characterized T2A-QF2_attp_3xp3-eYFP_attp donor backbone plasmid^4^, the 3xp3-eYFP sequence was removed using Xba1/Mlu1 restriction sites and the 3xp3-RFP coding sequence from pDSAR ^35^was amplified with gtagggtcgccgacatgacacaaggggtttctagaCGGTTCCCACAATGGTTAATTCG and cgacatgacacaaggggttctcgaGCCGTACCGTACGCGTATCGATAAGCTTTAAGATACATTGA before being inserted using SLiCE cloning ^36^.

Homology arms consisting of ~2kb immediately upstream and downstream of Cas9 cutsites were cloned into eYFP or RFP marked donor backbone plasmids using EcoRV/BamHI restriction sites for 5’ homology arms and Asc1/Spe1 for the 3’ homology arms. Homology arms were amplified from genomic DNA using the following PCR primer pairs:

IR93a pore-5HDR-fwd CCTGATCAACGAGGGTGAC

IR93a pore-5HDR-rev cgataccaGGATCCTATGTACCAGAAGCAGTTGTTCACC

IR93a pore-3HDR-fwd gtagcgaGGCGCGCCCTATGGTGCGCTACTGCAGC

IR93a pore-3HDR-rev TCGACactagtGGTTCATATGAGCGCTTTGG

IR93a TM2-5HDR-fwd CAGATATCGGACGAGGAGTG

IR93a TM2-5HDR-rev cgataccaGGATCCGGTACCGATGATGATTCGTCC

IR93a TM2-3HDR-fwd tagcaggcgcgccaTGGTGGTTGGGTATGTGCTG

IR93a TM2-3HDR-rev tcgacactagtGGTTCATATGAGCGCTTTGG

### Generation of transgenic mosquito strains

QUAS-GCaMP6f ^37^ and QUAS-*m*CD8:GFP ^38^ strains were a generous gift from Chris Potter. Generation of IR93a^pore^-T2A-QF2 and IR93a^TM2^-T2A-QF2 lines was accomplished using previously described methods^4^. Plasmids were prepared for injection using ZymoPURE II endotoxin-free midiprep kit (Zymo Research Corp, Irvine, CA). Cas9/sgRNA and donor plasmids were diluted to concentrations of 300ng/μl for injection. Freshly laid embryos were aligned against wet nitrocellulose paper and injected near the posterior pole using beveled aluminosilicate needles and a PLI-100 picoinjector (Harvard Apparatus). Surviving injected individuals were crossed to G3 stock and their F1 progeny were screened for fluorescence. Knock-in animals were further outcrossed to G3 for a minimum of 5 generations. To verify the correct insertion, animals were genotyped using a mix of three PCR primers (Universal forward primer 5’ – CCATCCTATGTGCGATCAACAA -3’, WT Rev 5’-GTACTTCGGTTCGGTCGAGTC-3’ and IR93a Rev 5’-CCGTATTGGCCACGTGTCC-3’). PCR products were run on an agarose gel where WT and mutant alleles could be clearly distinguished based on size (586 bp for WT, 261bp for IR93a^pore^, and 412 bp for IR93a^TM2^). The PCR products were further verified by Sanger sequencing.

### Single Sensillum Electrophysiology

Extracellular electrophysiological recordings from single antennal sensilla were performed as previously described^4^. Female mosquitoes (1-4 days old) were secured to glass slides with double sided tape and the antennae were flattened with human hair. A glass reference electrode was inserted into the eye and a recording electrode (filled with 130mM NaCl, 5mM KCl, 2mM MgCl_2_, 2mMCaCl_2_, 36mM D-sucrose, 5mM HEPES, pH 7.3) was inserted into one of the coeloconic sensilla found at the most distal tip of the 13^th^ antennal segment. Thermal stimuli were delivered via two alternate streams of dry air passing over the antennae. Temperatures of the air streams were calibrated to 25°C and 30°C by initially passing them through a common tube submerged in a warm water bath after which the streams were split and the warmer airstream was passed through insulated tubing further heated with a resistor (Emron E5CSV). Temperature at the antennae was recorded using an IT23-thermocouple (Physitemp) and a Fluke 80TK thermocouple module. Electrical signals detected in the sensilla were amplified using a TasteProbe DTP-02 (Syntech) and digitized with PowerLab 8/30. Data was band-pass filtered (100Hz-3000Hz) and acquired at 20k/s with LabChart 7 (ADInstruments). Spike thresholds were calculated by selecting the minimum interpeak value from spike amplitude histograms in Lab Chart 7. Spike rates were calculated as weighted average of Instantaneous Spike Frequencies across a 1 sec triangular window centered at the indicated time. Different time windows after onset were used to calculate average cooling and heating responses as the rise in weighted average spike frequency upon cooling was faster and sharper than its drop upon warming.

### Imaging and immunohistochemistry

To image CD8:GFP driven by IR93a, freshly dissected antennae of female mosquitoes were mounted in 80% glycerol/20% PBS and immediately imaged on an LSM 880 laser scanning confocal microscope (Zeiss). Immunostaining of cryosectioned mosquito antennae was performed as described^4^. Isolated mosquito heads (Genotype: AgIR93a^pore^-T2A-QF2/+; QUAS-CD8:GFP/+) were fixed for 30 min with ice cold 4% paraformaldehyde in PBS-T. Fixative was removed by washing with PBS before heads were cryoprotected overnight in 25% sucrose. After cryoprotection, antennae were dissected from heads and mounted in OCT compound (Tissue-Tek). Sections (14-18 μm) were dried to slides for 30 min before fixing with 4% paraformaldehyde for 15 min at room temperature. Fixative was removed by washing with PBS-T before blocking with 10% normal goat serum in PBS-T for 1 hour at room temperature. Slides were then incubated with primary antibody (chicken anti-GFP, GFP-1010, Aves Lab) diluted 1:1000 in 10% normal goat serum in PBS-T for 48 h at 4 C. Slides were washed 6 times for 20 minutes and incubated in secondary antibody (Alexa Fluor 488 goat anti-chicken-488, #A11039, Invitrogen) diluted 1:200 in 10% normal goat serum for 3 hours in the dark at room temperature. Slides were washed 6 times for 20 min in PBS-T and mounted in Vectashield with DAPI (Vector Laboratories).

Whole-mount immunohistochemistry was adapted from ^38^. Female mosquito heads were removed and digested for 1.5 hours at 37°C with 5U/mL chitinase (Sigma C6137) and 100U/mL chymotrypsin (Sigma CHY5S) resuspended in HEPES larval buffer (119mM NaCl, 48 mM KCl, 2mM CaCl_2_, 2mM MgCl_2_, 25mM HEPES, pH 7.5). Heads were quickly washed with ZnFa fixative (0.25% ZnCl2, 135mM NaCl, 1.2% sucrose, 0.03% Triton X-100, 2% PFA) before fixing for 24 hours at room temperature. After fixation, antennae were dissected in fresh HBS (150mM NaCl, 5mM KCl, 25mM sucrose, 10mM HEPES, 5mM CaCl2, 0.03% Triton X-100) then dehydrated for 1 hour in 80% methanol/20% DMSO. Antennae were then washed with 0.1M Tris pH 7.4, 0.03% Triton X-100 prior to blocking overnight at 4°C (1X PBS, 1% DMSO, 5% normal goat serum, 0.03% Triton X-100). Primary antibodies (mouse anti-*Apocrypta bakeri* Orco monoclonal antibody #15B2 (1:50 dilution) and chicken anti-GFP, GFP-1010, Aves Lab (1:500 dilution)) were diluted in blocking buffer and incubated with the tissue for 72 hours at 4°C. Primary antibodies were washed 5 times with 0.03% PBS-TritonX-100, 1% DMSO for 30 minutes followed by an overnight was at 4°C. The next day, antennae were incubated with secondary antibodies diluted in blocking buffer for 72 hours at 4°C (Cy5 goat anti-mouse and Alexa Flour488 goat anti-chicken (1:200)). Secondary antibodies were removed with the wash protocol used for primary antibodies. Antennae were then washed with PBS and mounted with Vectashield with DAPI.

### Calcium imaging

Transcuticular calcium imaging was performed based on a previous study ^37^. Animals were anesthetized on ice and their wings and legs were removed. Antennae were affixed to a glass slide with double sided tape and human hair. Images were acquired from the medioventral aspect of the second flagellomere using an Olympus BX51WI microscope fitted with an Olympus SLMPlan 50×/0.45 objective and a Hamamatsu Orca-R2 camera recording at 4 frames/sec. Hygrosensory stimuli were applied as described previously ^10^. A dry airstream (500ml/min) was connected to a solenoid valve that split the stream into two paths. The dry stream (~7% RH) was passed through an empty glass flask while the humid airstream (~90% RH) was bubbled through a flask filled with distilled water. During imaging, one airstream flowed for the first 5s before switching to the other stream for 10s after which it returned to the original stimulus for 5s. The starting condition (dry or humid) was alternated between individuals. Recordings were processed in ImageJ using Stackreg ^39^ to correct for movement. Baselines (F_0_) were calculated by taking the average of frames 7-24 preceding the first change in humidity. Moist and dry responses were quantified by taking the average of 10 consecutive frames where responses were found to be maximal across samples (frames 41-50 for DWD paradigm and frames 46-55 for the WDW paradigm).

### Humidity seeking Assay

Newly eclosed female mosquitoes were housed with males for 4-12 days. Before testing, females were separated into cages of 25-73 animals and starved overnight. Two hours before the start of experiments, water was removed and cages were allowed to acclimate to testing room conditions (22-24°C and 25-46% RH). Immediately following the acclimation period, animals were machine aspirated and released into the experimental chamber by gentle shaking to avoid the use of human breath. The experimental setup consisted of a polypropylene cage (30×30×30cm, BugDorm-1) with two rectangular 8-well dishes (12.8×8.55×1.5cm) situated in the middle of the floor. One tray was empty and the other was half filled with water. Both trays were covered with custom 3D-printed mesh covers that allow water vapor to escape but prevent animals from being able to contact the liquid with their proboscis. Once released into the testing chamber, a camera (B00UMX3HEG) situated above the cage recorded the animals movements throughout the 1hr assay. The number of animals contacting the mesh lids of each tray was counted at 1 min intervals. To verify the health and responsiveness of the animals, human breath was applied to the cage at the conclusion of the experiment and the number of animals that took flight during the subsequent 2 min interval was quantified.

### Heat seeking Assay

Heat seeking assays were performed as previously described^4^. A custom white plexiglass box (28×40×16cm) with transparent front viewing panel was used as the behavioral chamber. Two temperature-controlled Peltier elements (Custom Thermoelectric, LLC, Bishopville, MD) were mounted on the rear wall of the box, covered by white printer paper to match the surroundings. Likewise, a white gas diffusion pad was mounted on the ceiling to allow for introduction of CO_2_. The night before an experiment, 46-50 mated female mosquitoes (7-14 days old) were anesthetized on ice and placed into the heat seeking box. Animals were starved overnight with access to water-saturated cotton balls. Mosquitoes were maintained on an inverted light/dark cycle such that the lights were on during the overnight acclimation period and switched off 1 hour before the start of the assay. Environmental conditions in the behavior room were 25°C with a relative humidity of 70%. The experiments began when a randomly assigned Peltier unit was heated to 37°C while the other was set to 26°C. After 1-minute, a 20-second pulse of 4% CO_2_ (4% CO_2_, 21% O_2_, balance N_2_) was introduced through the diffuser pad. Mosquito activity was recorded using a front-mounted infrared camera (B00UMX3HEG) recording at 30 images/second. The number of animals on each Peltier was manually counted at 5 second intervals before being normalized for the CO_2_ takeoff response of each trial. CO_2_ takeoff was assessed by analyzing the individuals in view just prior to CO_2_ pulse and quantifying the percentage that took flight after the pulse. A heat-seeking index was then defined as the percent of mosquitoes landed on a Peltier divided by the percent CO_2_ takeoff of the trial. A single trial of G3 mosquitoes failed to activate in response to CO_2_ application and was therefore excluded from analysis.

### Host Approach Assay

Host approach assays were performed as previously described^4^. Female mosquitoes (4-10 days old) were separated from males and housed in cages of 41-76 animals (Bugdorm, 17.5×17.5×17.5cm). Mosquitoes were starved overnight and the next day the cages were removed from the incubator to acclimate to testing room conditions for 2 hours without water. To begin the experiment, 5 short human breaths were blown into the cage to activate host seeking behaviors. A 3mm high plastic mesh spacer was placed on the roof of the cage and the hand of a human volunteer was placed on the spacer such that the mosquitoes could approach near to the hand but could not make direct contact with their proboscis or other appendages. The same female volunteer was used for all assays. An iPhone XR camera was used to record the entire 5-minute period the hand was presented to the animals, followed by an additional minute after the hand was removed. The number of animals accumulated under the hand was quantified at 5 second intervals for the duration of the assay and divided by the total number of animals in the cage to give the percentage approaching the host. Individuals were categorized as flying, resting, or probing by selecting a specific frame and manually assigning a category to each individual based on their movements seen in the prior and subsequent frames.

### Blood feeding Assay

Blood feeding assays were performed as previously described^4^. 47-76 mated female mosquitoes (5-15 days old) were sexed on ice and starved overnight. The next day, cages were removed from the incubator to acclimate to testing room conditions (21-24°C, ~50% RH) for ≥1 hr. After acclimating, human blood warmed to 37°C was provided through the mesh roof of the cage via a collagen membrane feeding system (Hemotek, ltd., Blackburn, UK). Animals were allowed to feed for 20 minutes, after which they were collected on ice and squished onto white filter paper to check for evidence of a bloodmeal. Overall cage health was assessed by performing a host approach assay on the cages >1hr before blood feeding.

### Statistics

In all cases, n refers to independent biological replicates involving different animals or groups of animals. Shapiro-Wilk tests were used to assess the normality of all data sets (p≤0.05 rejected normal distribution). Parametric tests were performed on groups with normally distributed data. For single comparisons, a two-sided unpaired t-test was used. ANOVA with Tukey HSD post doc test was used for multiple comparisons (JMP11, SAS). If any data set did not exhibit a normal distribution, non-parametric tests were performed. Nonparametric tests consisted of either Wilcoxon (for single comparisons) or Kruskal-Wallis followed by a Steel-Dwass or Steel with control post hoc test for multiple comparisons (JMP11, SAS). In box plots, the box represents inter-quartile range (IQR), midline represents median, and whiskers extend to the lowest or highest data point that falls within 1.5 times the IQR from the box edges.

The Shapiro-Wilk test and t-test/ANOVA/Tukey HSD or Wilcoxon/Kruskal-Wallis/Steel with control/Steel-Dwass values obtained are listed below:

For the Moist Cell responses in Fig. 2: Response to moist air: *+/Ir93a^pore^*: W=0.947, p = 0.702; *Ir93a^pore^*: W = 0.956, p = 0.768; t-test, p < 0.0001. Response to dry air: *+/Ir93a^pore^*: W=0.970, p = 0.897; *Ir93a^pore^*: W = 0.884, p = 0.208; t-test, p = 0.006.

For the Dry Cell responses in Fig. 2: Response to moist air: *+/Ir93a^pore^*: W=0.947, p = 0.704;; *Ir93a^pore^*: W= 0.882, p = 0.198; t-test, p = 0.0002. Response to dry air: *+/Ir93a^pore^*: W= 0.920, p = 0.471; *Ir93a^pore^*: W = 0.695, p = 0.002; Wilcoxon: H = 10.54, p = 0.001.

Humidity seeking assays in Fig. 2: For (% wet, average from 51 to 60 min) – (% dry, average from 51 to 60 min): *wt*: W = 0.885, p = 0.083; *Ir93a^pore^*: W = 0.892, p = 0.284; *Ir93a^TM2^*: W = 0.794, p = 0.012; *Ir21a^EGFP^*: W = 0.765, p = 0.0408; Kruskal-Wallis: H = 22.08, df = 3, p < 0.0001. Steel with Control (vs wild type): wt vs. *Ir93a^pore^*, p = 0.0059; wt vs. *Ir93a^TM2^*, p = 0.0005, wt vs. *Ir21a^EGFP^*, p = 0.850.

For CO_2_ takeoff at 60 min in response to breath in Fig. 2: *wt*: W = 0.861, p = 0.040; *Ir93a^pore^*: W = 0.846, p = 0.112; *Ir93a^TM2^*: W = 0.962, p = 0.810; *Ir21a^EGFP^*: W = 0.975, p = 0.908; Kruskal-Wallis: H = 5.527, df = 3, p = 0.137.

For the 30°C to 25°C cooling responses in Fig. 3: *wt*: W= 0.742, p = 0.03; *Ir93a^TM2^*: W=0.704, p = 0.01; Wilcoxon: H = 6.82, df = 1, p = 0.009.

For the 25°C to 30°C warming responses in Fig. 3: *wt*: W= 0.880, p = 0.31; *Ir93a^TM2^*: W=0.930, p = 0.59; t-test, p = 0.0026.

For heat-seeking indexes in Fig. 3: *wt*: W= 0.996, p = .999; *Ir93a^pore^*: W=0.754, p = 0.009; *Ir93a^pore/TM2^*: W=0.827, p =0.042. Kruskal-Wallis Test: H = 14.55, df = 2, p=0.0007. Steel-Dwass test (p-value of each pairwise comparison): *wt* vs. *Ir93a^pore^*, p = 0.0061; *wt* vs. *Ir93a^pore/TM2^*, p = 0.0029; *Ir93a^pore^* vs. *Ir93a^pore/TM2^*, p = 0.5014.

For CO_2_ take-off in heat-seeking assays in Extended Data Fig. 3: *wt*: W= 0.888, p = .263; *Ir93a^pore^*: W=0.882, p = 0.199; *Ir93a^pore/TM2^*: W=0.838, p =0.055. ANOVA [F(2,21) = 3.00, p =0.07].

For hand-seeking in Fig. 4d: Accumulation beneath hand at 60 sec: *wt*: W = 0.955, p = 0.724; *Ir93a^pore^*: W = 0.734, p = 0.014; *Ir93a^TM2^:* W = 0.904, p = 0.468; Kruskal-Wallis, H = 1.61, df = 2, p=0.45.

For hand-seeking in Fig. 4e: Accumulation beneath hand (average from 240 to 300 sec): *wt*: W = 0.955, p = 0.724; *Ir93a^pore^:* W = 0.924, p = 0.503; *Ir93a^TM2^:* W = 0.968, p = 0.880; ANOVA [F(2,20) = 13.59, p =0.0002]. Tukey HSD alpha = 0.01.

For hand-seeking in Fig. 4f: Percentage in flight at 240s: *wt*: W = 0.862, p = 0.080; *Ir93a^pore^:* W = 0.953, p = 0.753; *Ir93a^TM2^:* W = 0.949, p = 0.729; ANOVA [F(2,20) = 8.84, p =0.0018]. Tukey HSD alpha = 0.05.

For blood feeding in Fig. 4h: *wt*: W = 0.918, p = 0.304; *Ir21a^+7bp^*: W = 0.923, p = 0.419; *Ir93a^pore^*: W = 0.688, p = 0.0003; *Ir93a^TM2^*: W = 0.846, p = 0.183; Kruskal-Wallis, H = 23.97, df = 3, p<0.0001. Steel-Dwass test (p-value of each pairwise comparison): *wt* vs. *Ir21a^+7bp^*, p = 0.665; *wt* vs. *Ir93a^pore^*, p = 0.0013; *wt* vs. *Ir93a^TM2^*, p = 0.0119; *Ir21a^+7bp^* vs. *Ir93a^pore^,* p = 0.0060; *Ir21a^+7bp^* vs. *Ir93a^TM2^*, p = 0.0175; *Ir93a^pore^* vs. *Ir93a^TM2^*, p = 0.5249.

For blood feeding in Extended Data Fig. 4: 37°C collagen, W = 0.918, p = 0.310; 26°C collagen, W = 0.783, p = 0.125; 37°C parafilm, W = 0.920, p = 0.742; 26°C collagen, not applicable. ANOVA [F(3,19 = 21.25, p <0.001]. Tukey HSD alpha = 0.05.

## Code availability

The source code for the Calcium Imaging analysis was previously reported in ref. 12 and is available at a text file at DOI: 10.7554/eLife.26654.010.

## Acknowledgements

We thank F. Catteruccia and C. Potter for mosquito strains and advice, G. Settles for Schlieren image, S. McIver for discussions, K. Menuz and C.Y. Su for protocols, V. Ruta for anti-Orco antisera, C. Zhu for assistance with husbandry and gradient measurements, A. Daniels for assistance with cloning, and L. Huang, E. Marder, S. McIver, M. Rosbash and members of the Garrity lab for comments on the manuscript. Funding was provided by National Institute of Allergy and Infectious Diseases grants R01 AI122802 (to PAG), R01 AI157194 (PAG), R21 AI140018 (PAG) and F31 AI133945 (CG), National Science Foundation grant IOS 1557781 (PAG), National Institute of Neurological Disorders and Stroke grant 2T32NS007292-31 (WJL) and Charles A. King Trust Postdoctoral Research Fellowship, Bank of America, N.A., Co-Trustees (WJL).

## Author contributions

W.J.L, G.B., E.C.C., R.G. and P.A.G. designed experiments. W.J.L, E.C.C. and R.G. performed husbandry, genetics and behavior. W.J.L performed molecular biology and gene targeting. W.J.L., R.T. and C.G performed immunohistochemistry. G.B. performed electrophysiology and calcium imaging. W.J.L., G.B., R.A. and P.A.G. performed data analysis. W.J.L. and P.A.G. created the figures and wrote the paper, with input from all authors.

## Additional Information

**Supplementary Information** (Movie 1 and Movie 2) is available for this paper.

Correspondence and requests for materials should be addressed to PAG.

---

**Extended Data Fig. 1:**
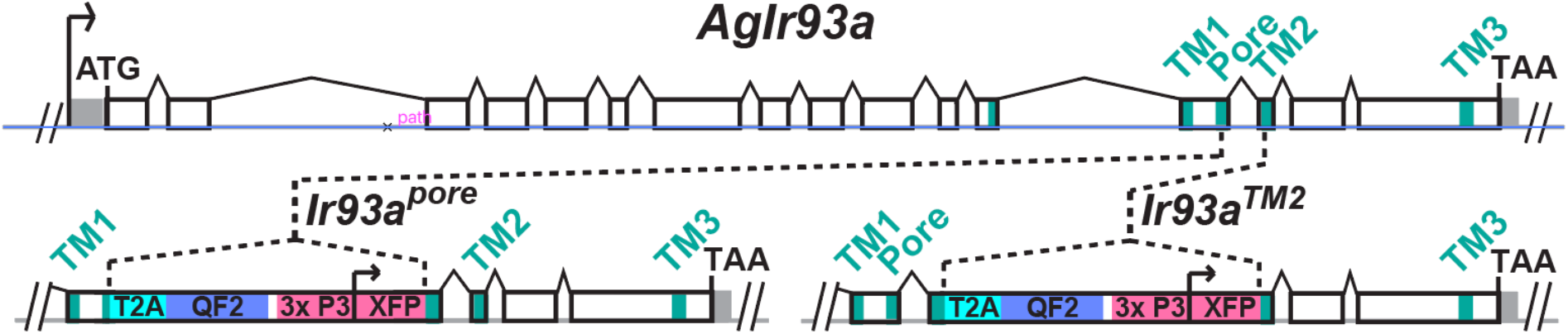
Organization of *Ir93a* locus in wild type and *Ir93a* knock-in animals. Top depicts *Ir93a* locus organization. Exonic regions encoding the three transmembrane domains and the putative ion pore are denoted in blue-green. 5’ and 3’ UTR are denoted in gray. Bottom depicts gene structure in *Ir93a^pore^* and in *Ir93a^TM2^*, each of which were produced by insertion of a knock-in cassette insertion at the gene location indicated. T2A sequence allows expression of two independent polypeptides from a single mRNA, permitting expression of the QF2 transcription factor as a separate polypeptide from a transcript initiated at the endogenous *Ir93a* start site. The 3xP3 promoter was used to drive retinal expression of the XFP fluorescent transformation marker. XFP was either RFP or YFP.

**Extended Data Fig. 2:**
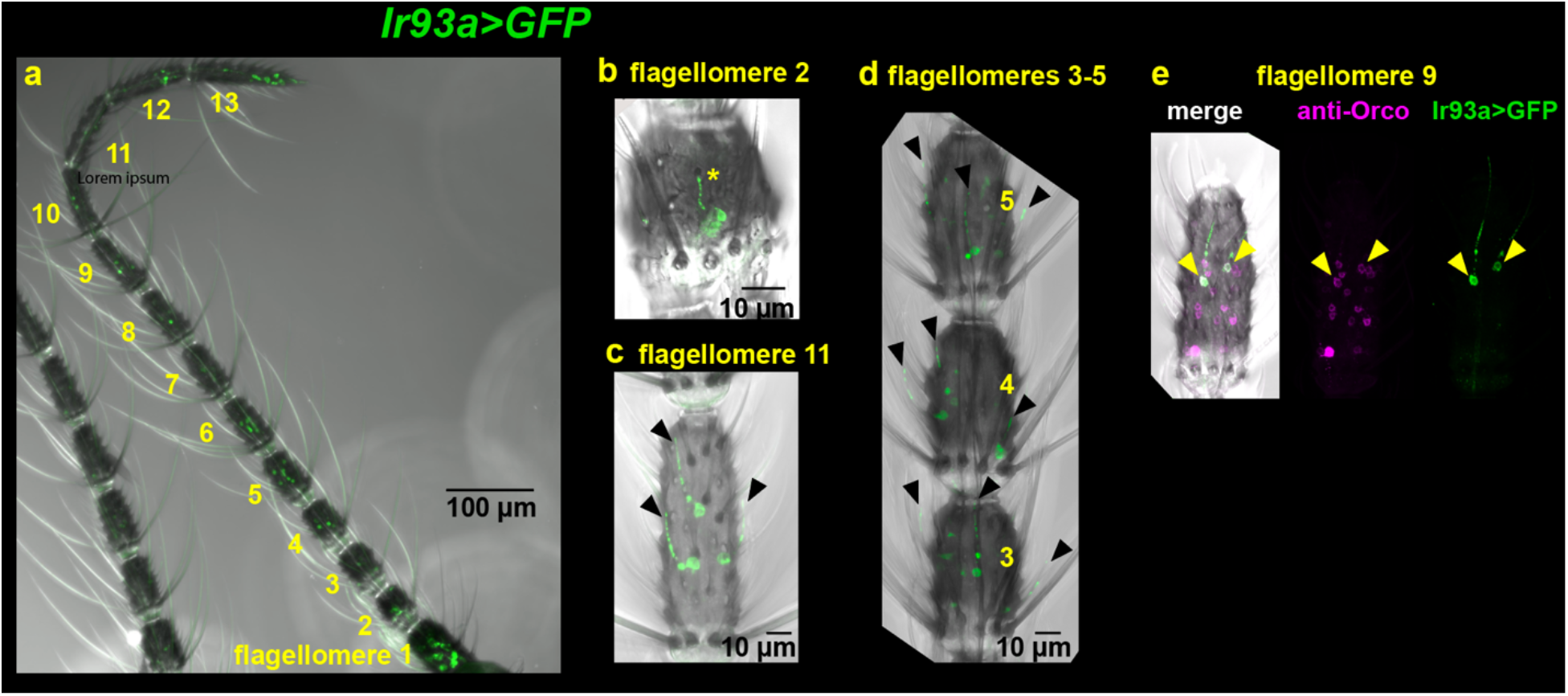
*Ir93a^pore^>mCD8:GFP* expression in mosquito antennae. **a,** *Ir93a>CD8:GFP* expression is detected in all 13 flagellomeres. **b,** Expression in neurons innervating sensilla ampullacea in flagellomere 2. **c-d**, Expression in neurons innervating trichoid sensilla in flagellomere 11 **(c)** and flagellomeres 3 to 5 **(d)**. Arrowheads mark GFP+ trichoid sensilla, asterisk marks entrance to sensilla ampullacea. **e**, *Ir93a>mCD8:GFP* expressing neurons that innervate trichoid sensilla co-express Orco.

**Extended Data Fig. 3:**
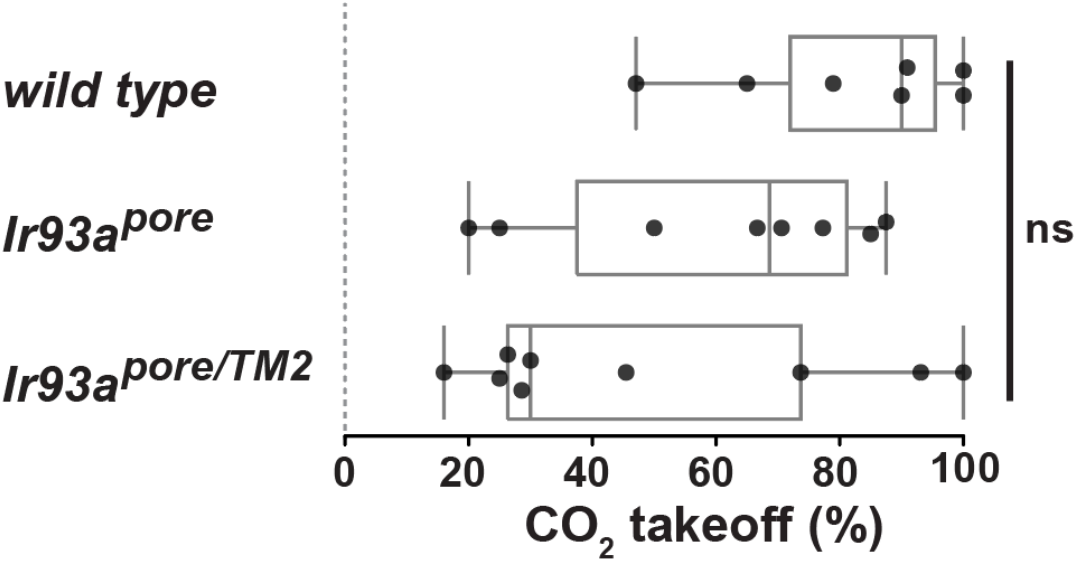
Responsiveness to CO_2_ in heat-seeking assay. Percentages of animals in heat-seeking assays that took flight after the initial pulse of 4% CO_2_.

**Extended Data Fig. 4.**
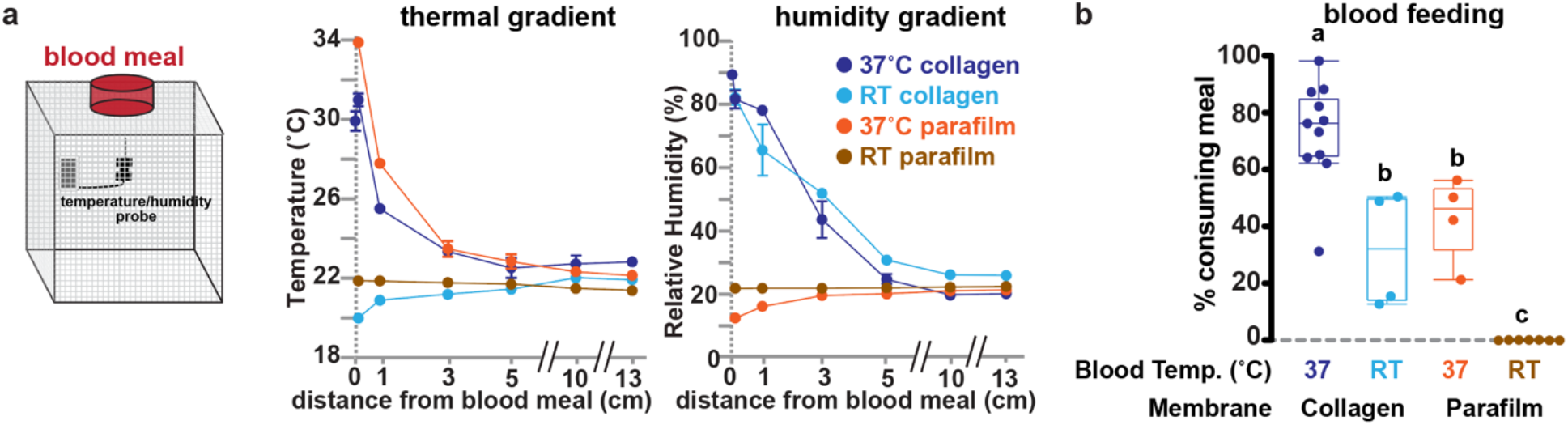
Using different membranes and blood meal temperatures induces different levels of blood feeding and creates different temperature and humidity gradients. **(a)** Left panel: Temperature and humidity probe was placed inside cage at multiple distances from blood meal. Center and Right panel: Temperature and relative humidity measurements at defined distances from blood meal presented using either a collagen (water vapor permeable) or parafilm (water vapor impermeable) feeding membrane. Ave. ± SEM. n = 3 measurements. Where SEM was smaller than dot, no error bar shown. **(b)** Blood feeding from meals at different temperatures and using different membranes. Letters denote distinct statistical groups, Tukey HSD, alpha = 0.05.

**Movie 1:** Human hand approach by G3 wild type mosquitoes activated by five breaths at start of assay. Movie is accelerated 8-fold.

**Movie 2:** Human hand approach by *Ir93a* mutant mosquitoes activated by five breaths at start of assay. Movie is accelerated 8-fold.

## Main References

1 World.Health.Organization. A global brief on vector-borne diseases. WHO/DCO/WHD/2014.1 (2014).

2 Lazzari, C. R. In the heat of the night. Science 367, 628–629, doi:10.1126/science.aba4484 (2020).

3 Carde, R. T. Multi-Cue Integration: How Female Mosquitoes Locate a Human Host. Current biology: CB 25, R793–795, doi:10.1016/j.cub.2015.07.057 (2015).

4 Greppi, C. et al. Mosquito heat seeking is driven by an ancestral cooling receptor. Science 367, 681–684, doi:10.1126/science.aay9847 (2020).

5 Benton, R., Vannice, K. S., Gomez-Diaz, C. & Vosshall, L. B. Variant ionotropic glutamate receptors as chemosensory receptors in Drosophila. Cell 136, 149–162 (2009).

6 Lewis, H. E., Foster, A. R., Mullan, B. J., Cox, R. N. & Clark, R. P. Aerodynamics of the human microenvironment. Lancet 1, 1273–1277, doi:10.1016/s0140-6736(69)92220-x (1969).

7 Craven, B. A. & Settles, G. S. A computational and experimental investigation of the human thermal plume. Journal of Fluids Engineering 128, 1251–1258 (2006).

8 Budelli, G. et al. Ionotropic Receptors Specify the Morphogenesis of Phasic Sensors Controlling Rapid Thermal Preference in Drosophila. Neuron 101, 738–747 e733, doi:10.1016/j.neuron.2018.12.022 (2019).

9 Knecht, Z. A. et al. Ionotropic Receptor-dependent moist and dry cells control hygrosensation in Drosophila. eLife 6, doi:10.7554/eLife.26654 (2017).

10 Knecht, Z. A. et al. Distinct combinations of variant ionotropic glutamate receptors mediate thermosensation and hygrosensation in Drosophila. eLife 5, doi:10.7554/eLife.17879 (2016).

11 Enjin, A. et al. Humidity sensing in Drosophila. Current biology: CB 26, 1352–1358, doi:10.1016/j.cub.2016.03.049 (2016).

12 Brady, O. J. et al. Vectorial capacity and vector control: reconsidering sensitivity to parameters for malaria elimination. Trans R Soc Trop Med Hyg 110, 107–117, doi:10.1093/trstmh/trv113 (2016).

13 Konopka, J. K. et al. Olfaction in Anopheles mosquitoes. Chem Senses 46, doi:10.1093/chemse/bjab021 (2021).

14 Brown, A. W. A., Sarkaria, D. S. & Thompson, R. P. Studies on the responses of the female Aedes mosquito. Part I. The search for attractant vapours. Bull Entomol Res 42, 105–114 (1951).

15 McIver, S. B. Sensilla mosquitoes (Diptera: Culicidae). J Med Entomol 19, 489–535 (1982).

16 Frank, D. D. et al. Early intergration of temperature and humidity stimuli in the Drosophila brain. Current Biology 27, 1–8, doi:10.1016/j.cub.2017.06.077 (2017).

17 Grimaldi, D. & Engel, M. S. Evolution of the insects. 489–491 (Cambridge University Press, 2005).

18 Pitts, R. J., Rinker, D. C., Jones, P. L., Rokas, A. & Zwiebel, L. J. Transcriptome profiling of chemosensory appendages in the malaria vector Anopheles gambiae reveals tissue- and sex-specific signatures of odor coding. BMC Genomics 12, 271, doi:10.1186/1471-2164-12-271 (2011).

19 Altner, H. & Loftus, R. Ultrastructure and function of insect thermo- and hygroreceptors. Ann. Rev. Entomol 30, 273–295 (1985).

20 Croset, V. et al. Ancient protostome origin of chemosensory ionotropic glutamate receptors and the evolution of insect taste and olfaction. PLoS Genet 6, e1001064, doi:10.1371/journal.pgen.1001064 (2010).

21 Eyun, S. I. et al. Evolutionary History of Chemosensory-Related Gene Families across the Arthropoda. Mol Biol Evol 34, 1838–1862, doi:10.1093/molbev/msx147 (2017).

22 Kozma, M. T. et al. Chemoreceptor proteins in the Caribbean spiny lobster, Panulirus argus: Expression of Ionotropic Receptors, Gustatory Receptors, and TRP channels in two chemosensory organs and brain. PloS one 13, e0203935, doi:10.1371/journal.pone.0203935 (2018).

23 Tichy, H. & Kallina, W. Insect hygroreceptor responses to continuous changes in humidity and air pressure. J Neurophysiol 103, 3274–3286, doi:10.1152/jn.01043.2009 (2010).

24 Chappuis, C. J., Beguin, S., Vlimant, M. & Guerin, P. M. Water vapour and heat combine to elicit biting and biting persistence in tsetse. Parasit Vectors 6, 240, doi:10.1186/1756-3305-6-240 (2013).

25 Gatehouse, A. G. The probing response of Stomoxys calcitrans to certain physical and olfactory stimuli. J. Insect Physiol. 16, 61–74 (1970).

26 Barrozo, R. B., Manrique, G. & Lazzari, C. R. The role of water vapour in the orientation behaviour of the blood-sucking bug Triatoma infestans (Hemiptera, Reduviidae). J Insect Physiol 49, 315–321, doi:10.1016/s0022-1910(03)00005-2 (2003).

27 Wiegmann, B. M. et al. Episodic radiations in the fly tree of life. Proceedings of the National Academy of Sciences of the United States of America 108, 5690–5695, doi:10.1073/pnas.1012675108 (2011).

28 Xiao, R. & Xu, X. Z. S. Temperature sensation: from molecular thermosensors to neural circuits and coding principles. Annual Review of Physiology 83, 205–230, doi:10.1146/annurev-physiol-031220-095215 (2021).

29 Zhao, Z. et al. Chemical signatures of human odour generate a unique neural code in the brain of Aedes aegypti mosquitoes. bioRxiv, DOI: 10.1101/2020.1111.1101.363861, doi:10.1101/2020.11.01.363861 (2020).

30 Littig, K. S. & Stovanovich, C. J. in Pictorial keys to arthropods, reptiles, birds and mammals of public health significance 134 (Communicable Disease Center, U.S. Department of Health, Education and Welfare, Public Health Service, 1966).

31 Giribet, G. & Edgecombe, G. D. The Phylogeny and Evolutionary History of Arthropods. Current biology: CB 29, R592–R602, doi:10.1016/j.cub.2019.04.057 (2019).

32 Penalver, E. et al. Ticks parasitised feathered dinosaurs as revealed by Cretaceous amber assemblages. Nature communications 9, 472, doi:10.1038/s41467-018-02913-w (2018).

33 Otálora-Luna, F., A. J. Pérez-Sánchez, Sandoval, C. & Aldana, E. Evolution of hematophagous habit in Triatominae (Heteroptera: Reduviidae). Revista Chilena de Historia Natural 88, 4, doi:10.1186/s40693-014-0032-0 (2015).

34 Rota-Stabelli, O., Daley, A. C. & Pisani, D. Molecular timetrees reveal a Cambrian colonization of land and a new scenario for ecdysozoan evolution. Current biology: CB 23, 392–398 (2013).

## Methods References

35 Volohonsky, G. et al. Tools for Anopheles gambiae Transgenesis. G3 (Bethesda) 5, 1151–1163, doi:10.1534/g3.115.016808 (2015).

36 Zhang, Y., Werling, U. & Edelmann, W. Seamless Ligation Cloning Extract (SLiCE) cloning method. Methods Mol Biol 1116, 235–244, doi:10.1007/978-1-62703-764-8_16 (2014).

37 Afify, A., Betz, J. F., Riabinina, O., Lahondere, C. & Potter, C. J. Commonly Used Insect Repellents Hide Human Odors from Anopheles Mosquitoes. Current biology: CB 29, 3669–3680 e3665, doi:10.1016/j.cub.2019.09.007 (2019).

38 Riabinina, O. et al. Organization of olfactory centres in the malaria mosquito Anopheles gambiae. Nature communications 7, 13010, doi:10.1038/ncomms13010 (2016).

39 Thevenaz, P., Ruttimann, U. E. & Unser, M. A pyramid approach to subpixel registration based on intensity. IEEE Trans Image Process 7, 27–41, doi:10.1109/83.650848 (1998).

